# Activation of Protein Kinase R (PKR) Plays a Pro-Viral Role in Mammarenavirus Infected Cells

**DOI:** 10.1101/2023.12.05.570143

**Authors:** Haydar Witwit, Roaa Khafaji, Arul Salaniwal, Arthur S. Kim, Beatrice Cubitt, Nathaniel Jackson, Chengjin Ye, Susan R Weiss, Luis Martinez-Sobrido, Juan Carlos de la Torre

**Affiliations:** Department of Immunology and Microbiology, The Scripps Research Institute, La Jolla, CA 92037; Department of Chemistry, The Scripps Research Institute, La Jolla, CA 92037; Texas Biomedical Research Institute, San Antonio, TX, USA; Department of Microbiology, Perelman School of Medicine at the University of Pennsylvania, Philadelphia, PA 19104

## Abstract

Many viruses, including mammarenaviruses, have evolved mechanisms to counteract different components of the host cell innate immunity, which is required to facilitate robust virus multiplication. The double strand (ds)RNA sensor protein kinase receptor (PKR) pathway plays a critical role in the cell antiviral response. Whether PKR can restrict the multiplication of the Old World mammarenavirus lymphocytic choriomeningitis virus (LCMV) and the mechanisms by which LCMV may counteract the antiviral functions of PKR have not yet been investigated. Here we present evidence that LCMV infection results in very limited levels of PKR activation, but LCMV multiplication is enhanced in the absence of PKR. In contrast, infection with a recombinant LCMV with a mutation affecting the 3’-5’ exonuclease (ExoN) activity of the viral nucleoprotein (NP) resulted in robust PKR activation in the absence of detectable levels of dsRNA, which was associated with severely restricted virus multiplication that was alleviated in the absence of PKR. However, pharmacological inhibition of PKR activation resulted in reduced levels of LCMV multiplication. These findings uncovered a complex role of the PKR pathway in LCMV-infected cells involving both pro-and anti- viral activities.

**IMPORTANCE:** As with many other viruses, the prototypic Old World mammarenavirus lymphocytic choriomeningitis virus (LCMV) can interfere with the host cell innate immune response to infection, which includes the double strand (ds)RNA sensor protein kinase receptor (PKR) pathway. A detailed understanding of LCMV-PKR interactions can provide novel insights about mammarenavirus-host cell interactions and facilitate the development of effective antiviral strategies against human pathogenic mammarenaviruses. In the present work, we present evidence that LCMV multiplication is enhanced in PKR-deficient cells, but pharmacological inhibition of PKR activation unexpectedly resulted in severely restricted propagation of LCMV. Likewise, we document a robust PKR activation in LCMV-infected cells in the absence of detectable levels of dsRNA. Our findings have revealed a complex role of the PKR pathway during LCMV infection and uncovered the activation of PKR as a druggable target for the development of antiviral drugs against human pathogenic mammarenaviruses.

## INTRODUCTION

Mammarenaviruses are enveloped viruses with a bi-segmented negative-stranded RNA genome (1), which cause chronic infections of their natural rodent host across the world, and human infections occur through mucosal exposure to aerosols or by direct contact of abraded skin with infectious materials (2). Upon a zoonotic event, mammarenaviruses can subvert the innate immune responses in the infected individual and interfere with the development of an effective adaptive immune response, which results in unrestricted virus multiplication and associated pathology and disease. Thus, several mammarenaviruses, chiefly Lassa virus (LASV), cause hemorrhagic fever disease in humans and pose important public health problems in their endemic regions (3). Moreover, evidence indicates that the worldwide distributed LCMV is a neglected human pathogen of clinical significance (4–8) and a serious risk to immunocompromised individuals (9, 10). There are no FDA-licensed arenavirus vaccines and current anti-arenaviral therapy is limited to an off-label use of ribavirin for which efficacy remains controversial (11). Therefore, there is a pressing need to develop effective strategies to combat human pathogenic mammarenaviruses, a task that will be facilitated by a better understanding of mammarenavirus-host innate defense interactions.

During multiplication in their host cells, RNA viruses generate a variety of pathogen-associated molecular patterns (PAMPs), including RNA-based PAMPs, that are recognized by host cellular pathogen recognition receptor (PRR) molecules, including TLR 3/7 and the retinoic acid-inducible gene I (RIG-I)-like receptors (RLRs) RIG-I and MDA5 (12, 13). Activated RLRs associate with the adapter mitochondrial antiviral-signaling (MAVS) protein (14) to promote activation of the non-classical IkB kinase (IKK)-related kinases (15) TANK-binding kinase 1 (TBK-1) and IkB kinase epsilon (IKKε) (16, 17) that activate interferon regulatory factor 3 (IRF3) and nuclear factor kappa-light- chain-enhancer of activated B cells (NF-κB), which together with ATF2/c-JUN initiate transcription of interferon beta (IFNβ) promoter (18). Interaction of secreted IFNβ with its receptor (IFNAR) activates the JAK/STAT signaling pathway (19) resulting in induction of hundreds of type 1 IFN (T1IFN) stimulated genes (ISGs) to produce a cellular antiviral state and control viral infection. Many viruses, including mammarenaviruses (20), have developed ways to subvert the T1IFN response (21, 22). Induction of IFNβ production in mammarenavirus-infected cells is greatly diminished by the virus nucleoprotein (NP) ability to inhibit activation of IRF3 (23, 24). The anti-IFNβ activity of NP has been linked to its C-terminally located functional 3’-5’ exonuclease (ExoN) domain characteristic of the DEDDh ExoN superfamily (25, 26). It is thought that NP’s ExoN activity promotes the degradation of viral RNA species that can activate the RIG- I/MAVS pathway and subsequent induction of IFNβ production to trigger the T1IFN pathway (27). The double strand (ds)RNA sensor protein kinase receptor (PKR) is at the center of cellular responses to a variety of stress signals, including viral infection (28). Short viral dsRNA species generated during virus replication in infected cells can trigger PKR activation reflected in increased levels of phosphorylated PKR (pPKR) (29–32). Increased levels of pPKR lead to phosphorylation of the eukaryotic translation initiation factor 2 alpha (eIF2α) resulting in the inhibition of cap-dependent protein translation initiation, which contributes to restricting virus propagation (33). Accordingly, many viruses have evolved mechanisms to counteract the activation of the PKR pathway (34–36). However, conflicting findings have been reported on mammarenavirus-PKR pathway interactions. The New World mammarenavirus Junin virus (JUNV) was reported to induce high levels of pPKR that did not result in increased levels of phosphorylated (p)eIF2α, whereas infection with the Old World mammarenavirus LCMV resulted in significantly increased levels of peIF2α despite minimal increased levels of pPKR (37). In addition, infection with the New World mammarenavirus Tacaribe virus (TCRV) resulted in increased levels of both pPKR and peIF2α, which contributed to restricted TCRV multiplication (38). However, other reports documented minimal or negligible activation of both PKR and/or eIF2α in cells infected with mammarenaviruses (39–42). Hence, the need for a detailed characterization of PKR-LCMV interactions is warranted, which is the focus of the present work.

Here, we investigated the interaction of LCMV with the PKR pathway and showed that LCMV infection resulted in minimal increased levels of pPKR, but LCMV multiplication was enhanced in PKR knock-out (PKR-KO) cells. However, unexpectedly, pharmacological inhibition of PKR activation resulted in severely restricted multiplication of LCMV, as well as LASV and JUNV. In contrast, cells infected with rLCMV/NP(D382A), impaired in its NP ExoN and anti-T1IFN activities (25), exhibited high levels of pPKR and peIF2α. However, levels of dsRNA remained below detectable levels in rLCMV/NP(D382A) infected cells. Our findings have uncovered a previously unnoticed proviral activity of the PKR pathway during mammarenavirus infection, raising the intriguing possibility that PKR activation might be considered as a potential druggable target to combat infections by human pathogenic mammarenaviruses.

## RESULTS

### Effect of genetic ablation of PKR on multiplication of LCMV

To determine whether genetic ablation of PKR affected LCMV multiplication, we compared the multi-step growth kinetics of LCMV/WT and the ExoN mutant rLCMV/NP(D382A) in WT and PKR knock-out (KO) A549 cells. We reasoned that LCMV/WT has already a high multiplication fitness in cultured cells, and therefore ablation of an antiviral host cell factor would be expected to result in only a modest increase in virus peak titers. In contrast, the fitness of an attenuated LCMV mutant, like rLCMV/NP(D382A), may be more dramatically affected by the removal of an antiviral host cell factor. We observed that LCMV/WT replicated to high titers in WT and KO A549 lines (Fig. 1A), but virus titers were consistently higher in PKR-KO compared to WT A549 cells. Multiplication of rLCMV/NP(D382A) was severely restricted in WT A549 cells and production of infectious progeny in TCS was only detected at 24 h pi and at very low levels (< 10^2^ FFU/mL), whereas at 48, 72 and 96 h pi titers in TCS were below detection levels (Fig. 1A). In contrast, rLCMV/NP(D382A) was able to multiply in PKR KO A549 cells resulting in titers of infectious progeny in TCS of >10^3^ FFU/mL (Fig 1A). We further confirmed the effect of PKR on LCMV multiplication in cultured cells by determining levels of LCMV NP RNA using RT-qPCR in WT and PKR-KO A549 infected cells (Fig. 1B). Consistent with the RT-qPCR results, levels of rLCMV/NP(D382A), but not LCMV/WT, small (S) segment RNA (replication) and NP mRNA (transcription) were drastically reduced in WT compared to PKR-KO A549 cells as determined by northern blotting (Fig. 1C). We next asked if the restricted multiplication of rLCMV/NP(D382A) in A549 WT cells could be overcome by increasing the initial virus input. We infected WT and PKR-KO A549 cells with either LCMV/WT or rLCMV/NP(D382A) using different MOIs and determined the numbers of NP^+^ cells at 48 h pi (Fig. 1D). Propagation of LCMV/WT was similarly efficient in WT and PKR-KO A549 cells, even at the lowest (0.05) MOI. In contrast, propagation of rLCMV/NP(D328A) in WT, but not PKR-KO, A549 cells was greatly influenced by the MOI used to initiate infection (Fig.1Di).

**Figure 1.**
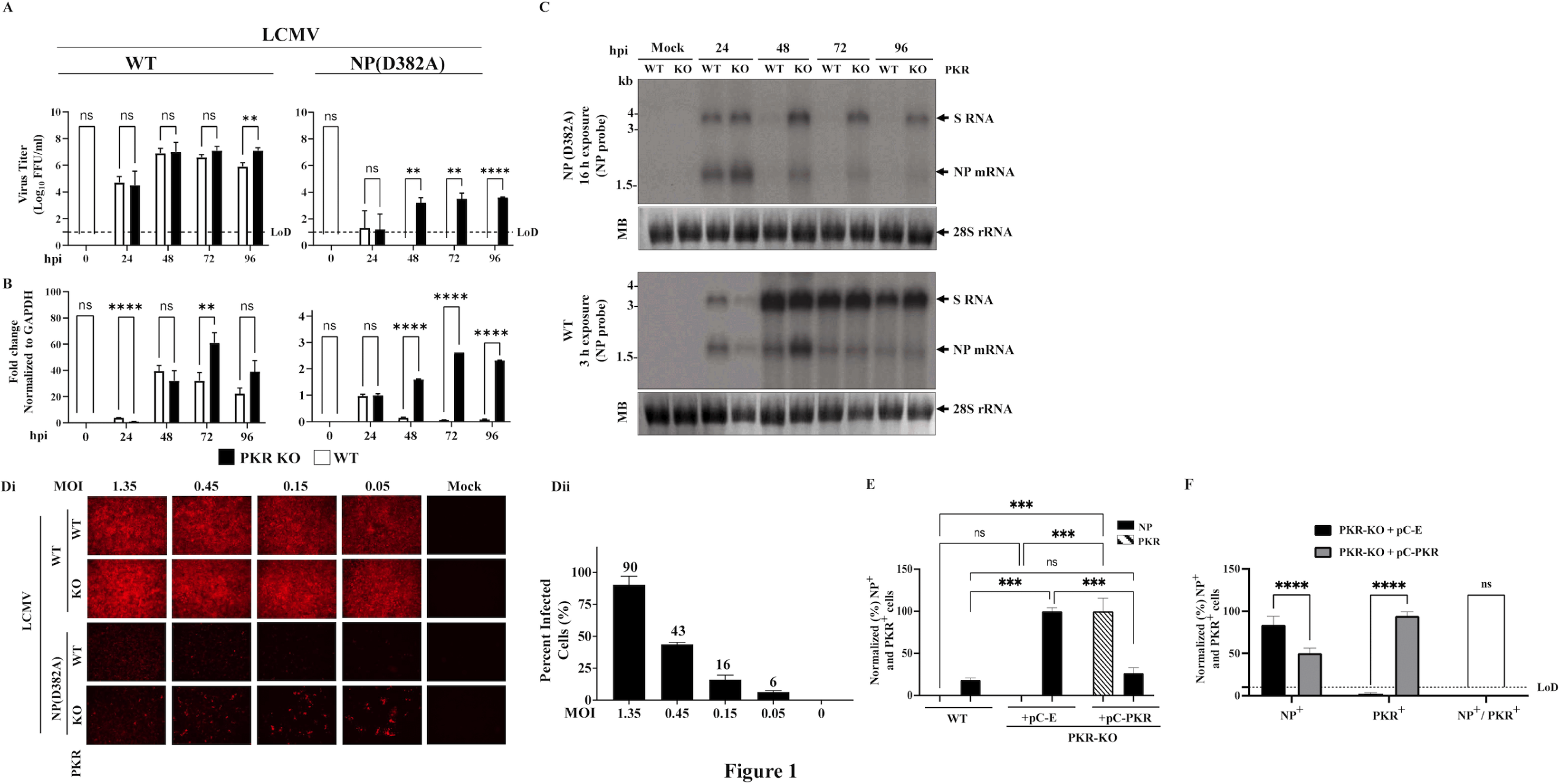
Multiplication of LCMV/WT and rLCMV/NP(D382A) in A549 cells. A. WT or PKR-KO A549 cells (biological triplicates) were infected with LCMV/WT (MOI 0.01) or rLCMV/NP(D382A) (MOI 0.05), and at the indicated h pi virus titers in TCS were determined. Values correspond to the mean ± SD. **B.** Determination of NP gene expression levels by RT-qPCR. Total cellular RNA from samples in A was isolated using Tri-Reagent and NP gene expression was determined by RT-qPCR. GAPDH was used for normalization utilizing the fold change (2^-ΔΔCt^) calculation and standardized using the 24 h pi value. Normalized data were plotted as mean ± SD (error bars) with a statistically significant set at *p*< 0.05. **C.** Northern blot analysis. RNA samples from B were analyzed by Northern blot. Methylene blue (MB) staining was used to confirm similar transfer efficiency for all RNA samples. Statistically significant values are labeled with 1, 2, 3, and 4 stars for P values less than 0.01, 0.001, and 0.0001 respectively. **D.** Effect of MOI on multiplication of rLCMV/NP(D382A). WT or PKR-KO A549 cells were seeded at 2x10^4^ cells /well in 96 well plates (two biological replicates, three technical replicates), and infected with LCMV WT or rLCMV/NP(D382A) at the indicated MOIs. At 48 h pi, cells were fixed and analyzed by IF. **Di.** LCMV infected cells were identified by IF using the rat monoclonal antibody VL4 to NP, followed by an anti-rat antibody conjugated to AlexaFluor 568. Individual images were obtained using Keyence BZ-X710 imaging system using 1 second exposure time with PlanApo_λ 10x 0.45/4.00mm objective lens. Files containing the labeled images were transferred to a laptop for processing the data using ImageJ. PowerPoint (2019 version) was used to compile and arrange the individual images. Adobe Illustrator was used to align the panels within the composite. **Dii.** Quantification of A549 WT cells infected with rLCMV/NP(D382A). Cell nuclei were stained with DAPI and the percentage of infected cells was determined based on the ratio (DAPI^+^+ NP^+^)/DAPI^+^. ImageJ software was used to quantify the signal, and GraphPad Prism software was used to blot the data. **E, F.** Effect of PKR complementation on rLCMV/NP(D382A) multiplication in PKR-KO A549 cells. **E.** PKR- KO A549 cells were transfected with pCAGGS empty (pC-E) or pCAGGS expressing PKR-P2A-mCherry (pC-PKR) in suspension, then seeded as indicated per plate format, and next day infected with rLCMV/NP(D382A) at MOI of 0.05. At 48 or 72 h pi cells were fixed for imaging analysis. Numbers (%) of infected cells were normalized to vehicle-treated infected cells. To ensure accurate comparisons, the signals of the virus (NP) were normalized to the pC-E conditions, which exhibited the highest signal intensity. Conversely, PKR was normalized to PKR transfected cells, as determined by the mCherry signal. This normalization approach was adopted due to the absence of mCherry signal in both the WT and pC-E conditions. Quantification of co-localization of NP and PKR signals as a function of Pearson’s coefficient, Spearman’s Rank coefficient and area overlap. These measures were obtained from running BIOP JaCoP script (44) using ImageJ software. A control test was done using DAPI images, copied, changed RUI to red and green, merged, and ran BIOP JaCoP to detect the colocalization, where the correlation showed 100%, then the images were split, one image was translated by transformation to variable degrees, remerged and re-ran the software to observe the correlation, the correlation was less depending on the degree of transformation and overlapping, samples were in duplicates. **F.** PKR-KO A549 cells were transfected with pC-E or pC-PKR in suspension, then seeded at 1x10^6^ cells/well into an M6 well plate and next day infected with rLCMV/NP(D382A) at MOI of 0.05. At 72 h pi single cell suspensions were prepared using Accutase, and after two washes with FACS buffer, cells were fixed with 4% PFA for 20 minutes, permeabilized, then probed with primary and secondary antibodies, followed by two washes after each antibody, resuspended in FACS buffer and analyzed using ZE5 analyzer (Bio-Rad). Quantification was done using FlowJo v10.9, then followed by statistical analysis using GraphPad Prism. Two- way ANOVA with Šidák correction for multiple comparisons was used.

Infection of WT A549 cells with rLCMV/NP(D382A) at MOI of 0.05 or 1.35 resulted in low (6%) and high (90%), respectively, percentage of NP^+^ cells (Fig.1Dii). These findings suggest that PKR exerts an antiviral role during LCMV infection that is counteracted by the ExoN and anti-T1IFN activities of the virus NP.

To assess whether the enhanced multiplication of rLCMV/NP(D382A) in PKR-KO A549 cells was mainly due to the absence of PKR, we transfected PKR-KO A549 cells with pCAGGS PKR-P2A-mCherrry (pC-PKR), a plasmid expressing PKR tagged with mCherry and containing the self-cleaving peptide (P2A) sequence (43) between the PKR and mCherry open reading frames. This allowed us to use mCherry expression as a surrogate of PKR expression in transfected cells. As a control, we used empty pCAGGS plasmid (pC-E). At 24 h post-transfection, cells were infected with rLCMV/NP(D382A), and at 48 and 72 h pi infected cells were identified by staining with a NP-specific antibody. We observed a reduced level of rLCMV/NP(D382A) infection at 48 and 72 h pi in A549 PKR-KO cells expressing pC-PKR (mCherry+) compared to cells transfected with the control pC-E plasmid (Fig. 1E), which correlated with greatly reduced colocalization of NP (LCMV infection) and mCherry (PKR expression) as determined by Pearson’s coefficient, Spearman’s Rank coefficient and area overlap measures obtained from running BIOP JaCoP script (44) using ImageJ software. We further validated these findings by flow cytometry (Fig. 1F).

We also examined the role of other IFN-induced genes including RNase L and MAVS during LCMV infection, and found that MAVS, but not RNAse L, also contributed to the restricted multiplication of rLCMV/NP(D3282A) (data not shown).

### Effect of LCMV infection on PKR activation

Activation of PKR requires its autophosphorylation upon binding to viral dsRNA and phosphorylated PKR (pPKR) negatively regulates translation via phosphorylation of eIF2a. We found that rLCMV/NP(D382A), but not LCMV/WT, activated PKR as determined by the detection of increased levels of pPKR (Fig. 2A) and peIF2α (Fig. 2B). We used infection with Sindbis virus (SINV) as a positive control, as SINV infection activates the PKR pathway (31, 45). Quantification of the immunoblots (IB) showed that levels of pPKR were 4-fold higher in rLCMV/NP(D382A) than LCMV/WT infected cells, both normalized to levels of pPKR detected in mock-infected control cells (Fig. 2Aii).

**Figure 2.**
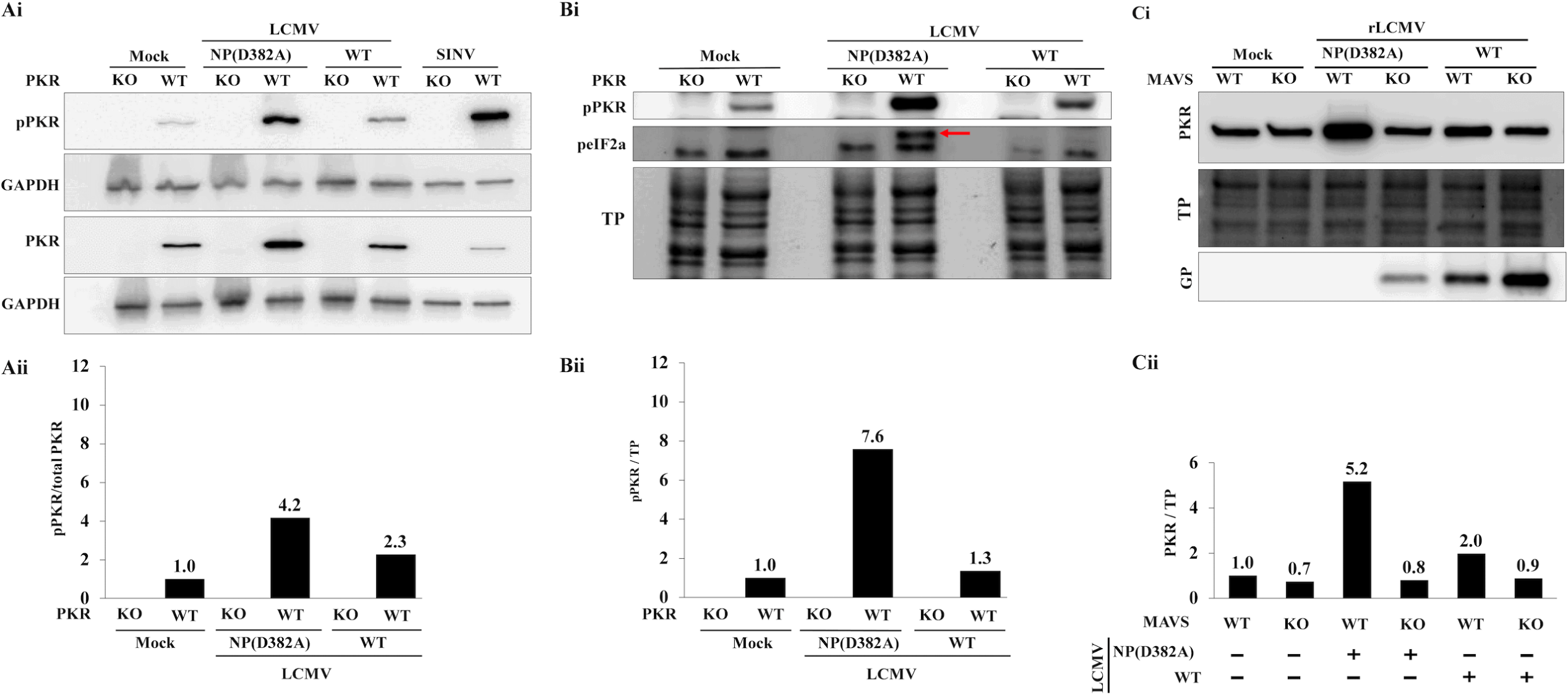
A. PKR activation in LCMV infected cells. Ai. WT or PKR-KO A549 cells were infected with LCMV WT (MOI 0.01), rLCMV/NP(D382A) (MOI 0.05), or SINV (MOI 3), and at 48 h pi cells lysates prepared for Western blot using the indicated antibodies. Samples from SINV-infected cells containing 8-fold lower amount of protein, compared to the other lysates, were used as a control (one-eighth). Levels of GAPDH were used for the normalization of samples. Membranes were probed with anti-PKR, anti-pPKR, or anti-GAPDH (primary), anti-GP and horseshoe peroxidase (secondary) antibodies. **Aii.** Bands of IB membranes were quantified, and the value of the ratio of pPKR/PKR plotted. **Bi.** WT and PKR-KO A549 cells were infected with LCMV WT (MOI 0.01) or rLCMV/NNP(D382) (MOI 0.05). At 48 h pi cell lysates were prepared for Western blot analysis. IB membrane was probed with anti-PKR, anti-pPKR, or anti-phospho-eIF2α (primary) and the corresponding HRP-conjugated secondary antibodies. Arrow refers to the eIF2α band. **Bii.** pPKR/total protein signal ratio. IB signals were quantified using Image Lab software (BioRad). Total protein values were used to normalize the values and normalized results were plotted. **C.** Interferon-dependent PKR-induced expression. **Ci.** WT and MAVS-KO A549 cells were seeded in a 6 well plate at a density of 1x10^6^ cells/well and infected with LCMV WT (MOI 0.01) or rLCMV/NP(D382A) (MOI 0.05). At 48 h pi, cell lysates were prepared for western blot analysis using anti-PKR and LCMV- GP antibodies. **Cii.** IB signals were quantified using Image Lab and normalized to the total protein signal.

Using total protein staining to normalize the signal in the IBs, we observed a higher level of pPKR, and significantly increased levels of peIF2α in cells infected with rLCMV/NP(D382A) but not in cells infected with LCMV/WT (Fig 2Bi and 2Bii).

PKR is an interferon-stimulated gene (ISG) and LCMV with mutations affecting NP’s ExoN activity (e.g. D382A) have been shown to induce much higher levels of T1IFN than LCMV/WT in infected cells, a process mediated by the cytosolic PRR RIG-I and subsequent activation of the MAVS adaptor and downstream activation of IRF3, a transcriptional factor that promotes induction of IFN-*β* expression (46). To assess whether the observed higher levels of total PKR protein in rLCMV/NP(D382A)-infected A549 cells was driven by the T1IFN pathway, we compared levels of PKR protein in WT and MAVS-KO A549 cells infected with LCMV/WT or rLCMV/NP(D382A) (Fig. 2C).

Levels of PKR were higher in rLCMV/NP(D382A) than LCMV/WT infected WT A549 cells, whereas both LCMV/WT and rLCMV/NP(D382A) infected MAVS-KO A549 cells exhibited similar levels of total PKR protein. Consistent with previous results (Fig.1), levels of LCMV GP2, as determined by western blot, were slightly higher in MAVS-KO compared to WT-infected A549 cells, whereas infection with rLCMV/NP(D382A) resulted in detectable levels of GP2 expression in MAVS-KO but not in WT A549 cells. These findings supported that T1IFN played a role in driving the expression of total PKR protein in rLCMV/NP(D328A) infected cells.

We next investigated whether the extremely low increased levels of pPKR in LCMV/WT infected cells reflected LCMV’s ability to either escape sensing by PKR or actively interfere with the activation of PKR. For this, we infected WT A549 cells with LCMV/WT (MOI 0.5) for 24 h and then superinfected them with SINV (MOI 3). At 24 h pi with SINV, we prepared whole cell lysates and analyzed them by Western blot (Fig. 3). SINV-mediated activation of PKR, as determined by increased levels of pPKR, was not significantly affected by LCMV infection, indicating that LCMV does not actively inhibit the activation of PKR. SINV infection did not affect LCMV multiplication in A549 cells based on levels of GP2 expression determined western blot (Fig. 3A and 3B) and numbers of LCMV NP+ cells determined by IF (Fig. 3C).

**Figure 3.**
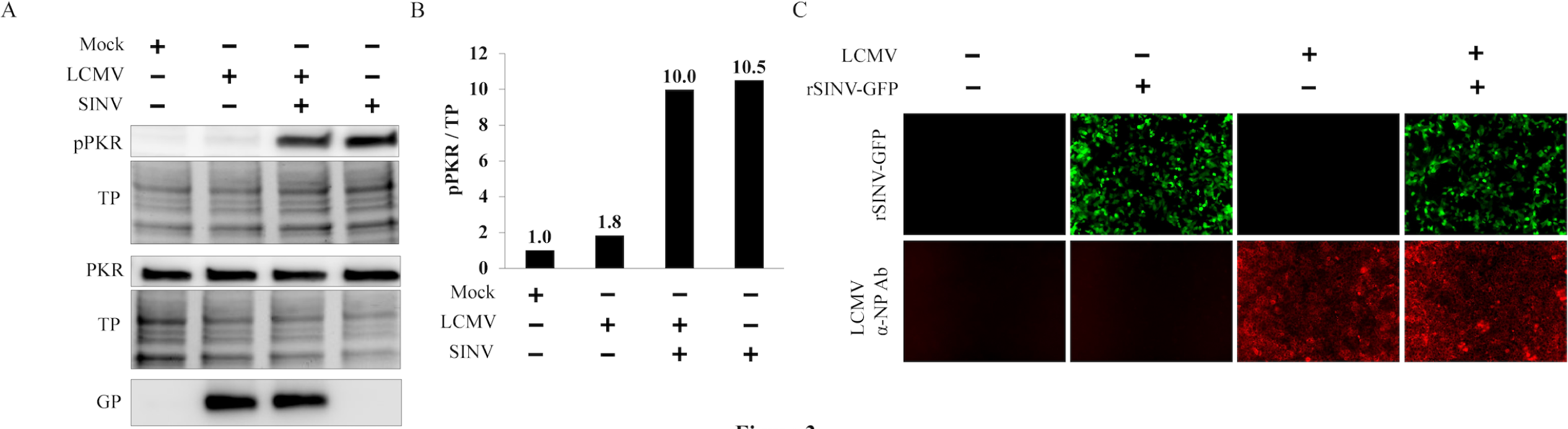
Effect of LCMV infection on PKR activation. **A.** WT A549 cells were infected with LCMV/WT (MOI 0.5) for 24 h and then superinfected with SINV (MOI 3). At 24 h pi with SINV, whole cell lysates were prepared for Western blot analysis using antibodies to PKR, pPKR or LCMV-GP. **B.** The IB signals were quantified using Image Lab and normalized to total protein signal. **C.** WT A549 cells were infected with LCMV/WT (MOI 0.5) for 24 h and then superinfected with SINV (MOI 3). At 24 h pi with SINV, cells were fixed for IF analysis using antibodies against LCMV-NP, images were taken at 20X magnification with Keyence BZ-X710.

### Role of dsRNA on PKR activation in rLCMV/NP(D382A)-infected cells

We next investigated whether increased levels of dsRNA correlated with increased levels of pPKR observed in rLCMV/NP(D382A) compared to LCMV/WT infected cells.

For this, we infected A549 WT cells with LCMV/WT or rLCMV/NP(D382A) for 24 h, or SINV as a positive control, and used the anti-dsRNA 9D5 antibody to detect dsRNA by immunofluorescence (IF). Both LCMV/WT and rLCMV/NP(D382A) infected cells, as well as mock-infected control cells, had undetectable levels of dsRNA, whereas dsRNA was readily detected in SINV-infected cells (Fig. 4A). We obtained similar results in IFN-deficient Vero E6 cells (Fig. 4B). Levels of dsRNA remained undetectable in Vero E6 cells that had been persistently infected with LCMV for 15 days, but SINV infection of LCMV persistently infected cells resulted in high levels of dsRNA detected by IF (Fig. 4B). Unexpectedly, in contrast to published findings (40), we did not detect dsRNA in cells infected with the Candid#1 live-attenuated strain of the New World Junin virus (JUNV). We obtained similar results using the anti-dsRNA J2 antibody (data not shown). These findings indicated that under our experimental conditions infection with either WT or NP(D382A) mutant LCMV does not result in levels of dsRNA that can be detected with the validated 9D5 antibody to dsRNA (47, 48).

**Figure 4.**
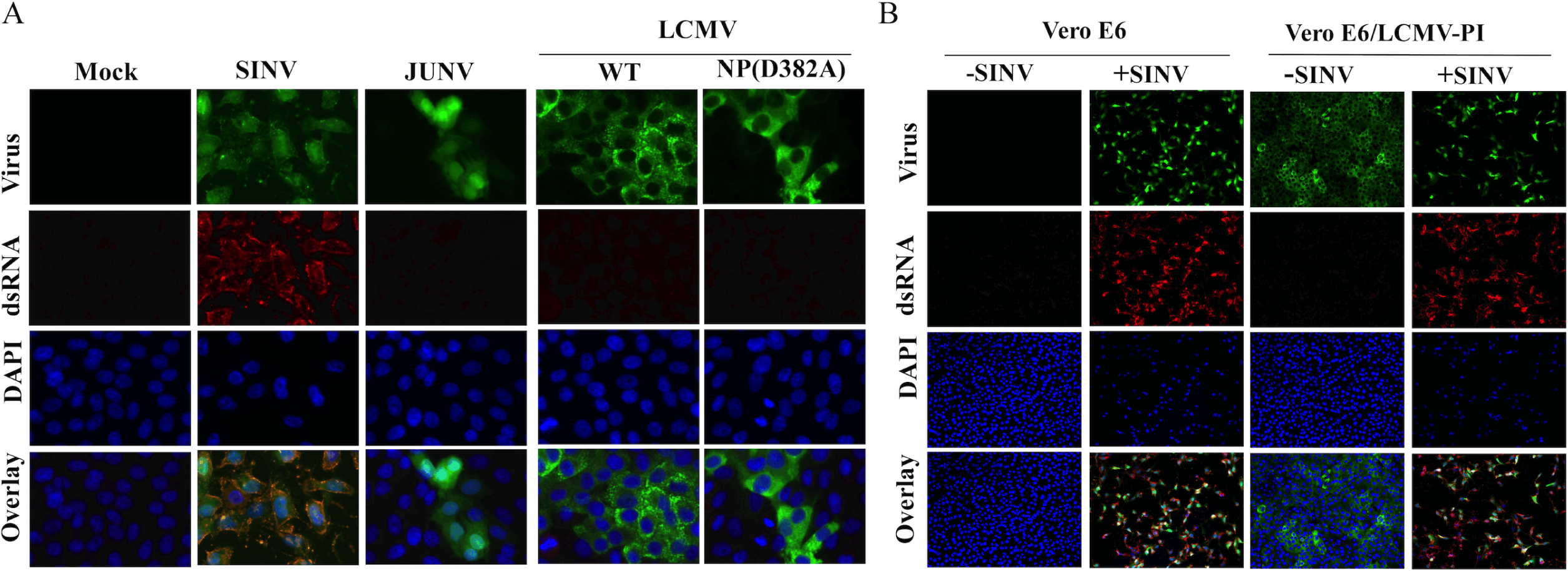
Detection of dsRNA. **A.** A549 cells were seeded at 1.5x10^4^ cells/well in a 96 well plate and infected with LCMV/WT or rLCMV/NP(D382A) at MOI of 0.5. At 24 h pi, cells were fixed and analyzed by IF using 9D5 anti-dsRNA antibody. Infection with SINV (MOI 3) was used as a positive control. **B.** IFN deficient Vero E6 cells were infected with LCMV/WT (MOI 0.01), rLCMV/NP(D382A) (MOI 0.05), Candid#1 live-attenuated vaccine strain of JUNV (MOI 0.5), and SINV (MOI 3). At 24 h pi cells were fixed and analyzed by IF as in A. Images were taken at 20X magnification and zoomed in digitally twice. **C.** Effect of LCMV persistence on the production of dsRNA. Vero E6 (left) or BHK21 (right) cells were infected with LCMV WT at MOI of 0.01 in a 6-well plate, then expanded to T25 and passaged every 3-4 days along parenteral mock-infected cells. On day 15, cells were seeded into a 96 well plate at a density of 2x10^4^ cells/well and incubated for 24 h, then superinfected with SINV (MOI 3). At 24 h pi with SINV, cells were fixed and processed for IF as in A. Images were taken using Keyence BZ-X710.

### Assessing the contribution of T1IFN to PKR activation in rLCMV/NP(D382A)- infected cells

We have shown that rLCMV/NP(D382A), but not LCMV/WT, can trigger a robust T1IFN response (25). The highly enhanced multiplication of rLCMV/NP(D382A) in PKR-KO compared to WT A549 cells could reflect differences in induction of T1IFN expression between WT and PKR-KO A549 cells following infection with rLCMV/NP(D382A). To examine this possibility, we infected WT or PKR-KO A549 cells with LCMV/WT (MOI 0.01) or rLCMV/NP(D382A) (MOI 0.05) and at 72 h pi collected TCS samples that we used to treat IFN deficient Vero E6 cells for 24 h, followed by their infection (MOI 0.1) with vesicular stomatitis virus (VSV). At 24 h pi with VSV we fixed the cells and stained them with crystal violet to assess VSV induced cytopathic effect (CPE) (Fig. 5A). Vero E6 cells treated with TCS from either WT or PKR-KO A549 cells infected with rLCMV/NP(D382A) were protected against VSV-induced CPE. As expected, Vero E6 cells treated with TCS from WT or PKR-KO A549 cells infected with LCMV/WT were fully susceptible to VSV induced CPE. Results of quantitative analysis of expression of the ISGs MX1 and ISG15 in Vero E6 cells treated with TCS from A549 cells infected with LCMV/WT or rLCMV/NP(D382A) (Fig. 5B) were consistent with the results of the T1IFN bioassay (Fig. 7A). ISG15 and MX1 genes showed a significant upregulation, ∼4 and ∼500 folds respectively, in Vero E6 cells treated with TCS from A549 cells infected with NP(D382A), but not LCMV/WT, compared to treatment with TCS from mock-infected A549 cells.

**Figure 5.**
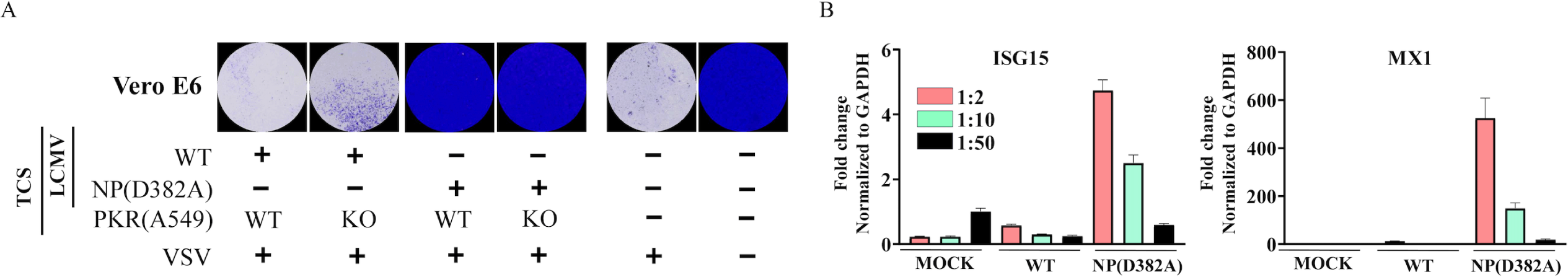
T1IFN response in PKR-KO A549 cells infected with LCMV. A. WT and PKR-KO A549 cells were seeded at 1x10^5^ cells/well in 24 well plate, and next day infected with LCMV/WT (MOI 0.01) or rLCMV/NP(D382A) (MOI 0.05). At 24 h pi, TCS were collected and used to treat Vero E6 cells seeded at 1x10^5^ cells/well in 24 well plate. After 24 h exposure to TCS, Vero E6 cells were infected with VSV at MOI of 0.1 and at 24 h pi with VSV, cells were fixed and stained with crystal violet for 10 minutes and imaged with BioSpot machine. **B.** TCS collected from 48 h pi of A549 WT infected with LCMV/WT or rLCMV/NP(D382A) were used to treat Vero E6 cells seeded at 1x10^5^ cells/well in 24 well plate. After 24 h exposure to TCS, cells were lysed with Tri-Reagent and RNA was collected and pretreated with ezDNAse for 5 min at 37°C, then 10ng/sample (three technical replicates) for RT-qPCR (two steps: RT with random hexamers, followed by qPCR with specific primers), Sybr green was used for RT-qPCR.

### Effect of pharmacological inhibition of PKR activation on LCMV multiplication

To assess the effect of PKR inhibition on LCMV multiplication, we examined the effect of the PKR inhibitor C16, and its inactive control C22, (49) on LCMV multiplication. Dose-response assays indicated that C16, but not C22, exerted a dose-dependent inhibitory effect on LCMV multiplication (Fig. 6A). C16 exhibited an EC_50_ = 0.177 µM and CC_50_ > 20 µM, while C22 exhibited an EC_50_ > 20 µM and CC_50_ > 20 µM (Fig. 6B). Treatment (3 µM) with C16, but not with C22, strongly inhibited propagation of LCMV in A540 cells following infection at low (0.01) MOI (Fig. 6C). Treatment with C16 (3 µM) caused > 5 logs reduction in peak titers of rLCMV/GFP in WT A549 cells (Fig. 6D). A much lower (∼ 2 logs) reduction in peak titers of rLCMV/GFP was observed in PKR-KO A549 cells treated with C16, which may reflect C16 off-target effects that affected PKR-independent pathways and factors (50).

**Figure 6.**
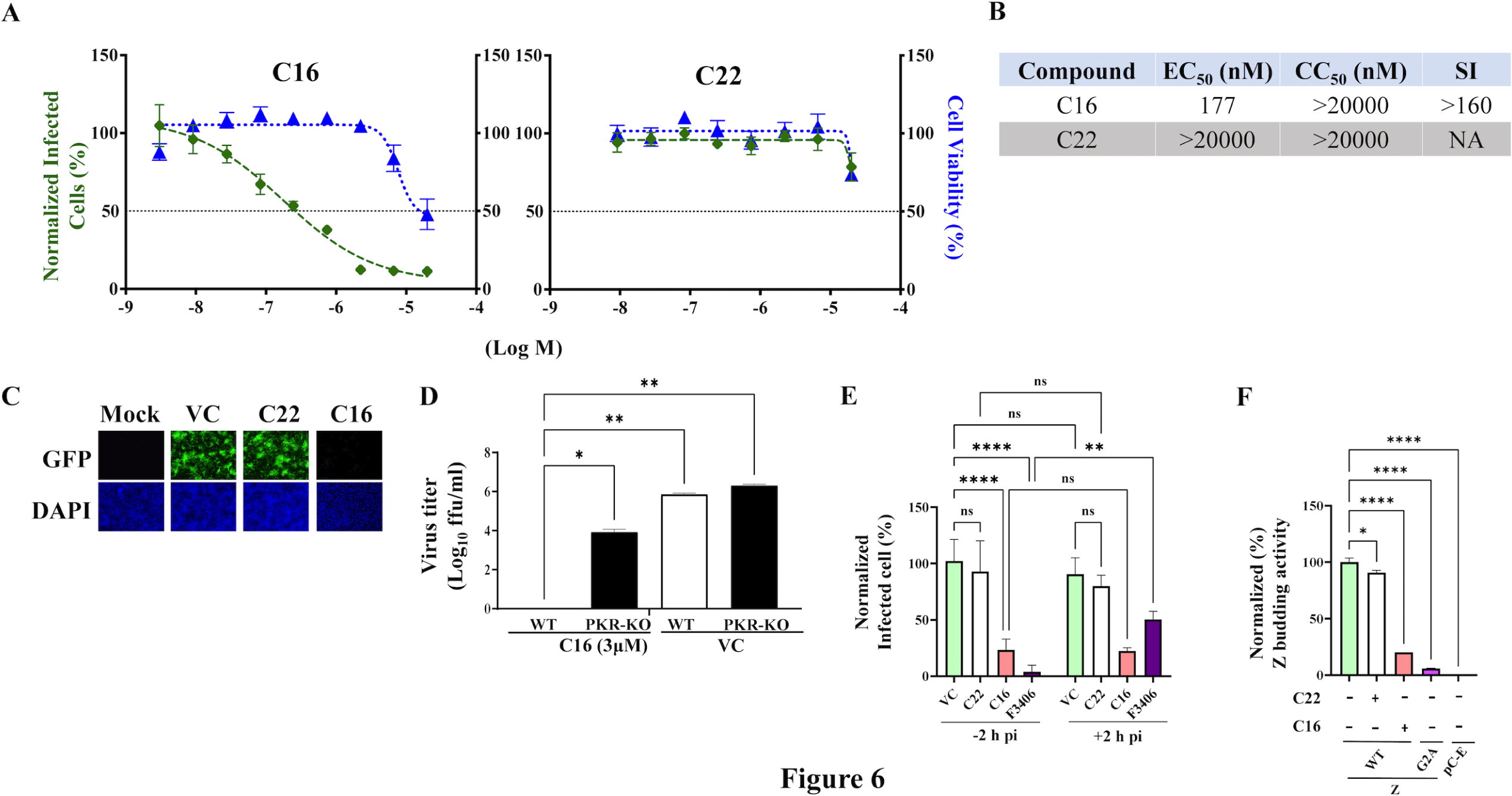
Effect of PKR inhibition on LCMV multiplication. A. C22 and C16 Dose- response. A549 cells were seeded at 2x10^4^ cells/well into a 96-well plate, infected with LCMV/WT (MOI of 0.05) and treated with C16 and C22 at the indicated concentrations. At 96 h pi cells were fixed and the numbers of infected cells were determined by IF using a rat mAb to NP. Numbers (%) of infected cells were normalized to vehicle-control infected cells. Results show the percentage of infected WT vs PKR-KO A549 cells (four replicates). **B.** C16 and C22 EC_50_, CC_50,_ and selectivity index (SI). **C.** Effect of C16 and C22 on LCMV propagation. A549 cells were seeded at 4x10^4^ cells/well into a 96-well plate, infected with LCMV (MOI of 0.05) and treated (3 µM) with either CC16, C22, or vehicle control (VC). At 96 h pi cells were fixed and stained with DAPI, IF analysis was determined using Keyence BZ-X710, images were taken at 10X magnification. **D.** Effect of C16 and C22 on the production of virus infectious progeny. A549 cells were seeded at 2x10^5^ cells/well in an M12-well plate, infected with LCMV/WT (MOI of 0.01) and treated (3 µM) with C16 or C22 compounds, or with VC. At 72 h pi, TCS were collected, and virus titers were determined by FFA using Vero E6 cells in a 96-well plate format. The repeated measures ANOVA with mixed effect analysis, and Dunnet correction for multiple comparisons were used. **E.** Time of Addition assay. A549 cells were seeded into a 96 well plate at a density of 2 x 10^4^ /well. Next day, cells were infected with the single-cycle infectious rLCMVΔGPC/ZsG (MOI = 0.5) and treated (3 µM) with C16 or C22 compounds, or with VC, starting 2 hours prior (- 2 h) or after (+ 2 h) infection. The LCMV cell entry inhibitor F3406 (5 μM) was used as a control. At 48 h pi, ZsG^+^ cells were assessed using Cytation 5 reader, values were normalized to vehicle control treated and infected cells. **F. Budding Assay.** HEK293T cells were seeded at a density of on poly-l-lysine coated wells in M12 well format. Next day, cells were transfected with either pC.LASV-Z-GLuc or pC.LASV-Z-G2A-GLuc (mutant control) or pCAGGS- Empty(pC-E. At 5 h post transfection, cells were washed three times and fed with fresh medium containing the indicated drugs and concentrations. At 48 h post-transfection both TCS were collected and whole cell lysis (WCL) prepared. GLuc activity was determined in TCS and WCL using SteadyGlo Luciferase Pierce: Gaussia Luciferase Glow assay kit utilizing Cytation5 reader, raw data signal was normalized and then plotted using GraphPad Prism software (v10).

To gain insights about the mechanism by which the PKR kinase inhibitor C16 disrupted the LCMV life cycle, we used C16, and its negative control C22, in cell-based assays for different steps of the LCMV life cycle. To distinguish between an effect of C16 on a cell entry or post-entry step of LCMV, we conducted a time of addition assay using a single cycle infectious recombinant LCMV expressing GFP (rLCMVΔGPC/GFP) (51). C16 inhibited multiplication of rLCMVΔGPC/GFP, whether applied 2 hours before (-2 hpi) or 2 hours after (+2 hpi) infection (Fig. 6E). In contrast, regardless of the time of its addition, C22 did not inhibit rLCMVΔGPC/GFP. As predicted addition at -2 hpi, but not at +2 hpi, of the LCMV entry inhibitor F3406 (52), resulted in inhibition of rLCMVΔGPC/GFP. This observation indicated that C16 inhibited a post cell entry step of the LCMV life cycle. To assess the effect of C16 on mammarenavirus budding, a process driven by the viral matrix protein Z (53), we examined the effect of C16 on the budding activity of the LCMV matrix Z protein using a published cell-based assay where levels of Gaussia luciferase (GLuc) activity serve as an accurate surrogate of Z expression levels (54). C16, but not C22, inhibited Z budding activity (Fig. 6F).

### Effect of pharmacological inhibition of PKR on multiplication of other mammarenaviruses

To determine whether our findings could be also extended to other mammarenaviruses, we examined whether multiplication of the New World JUNV and Old World LASV mammarenaviruses were also affected by pharmacological inhibition of PKR activation. For this, we infected A549 WT cells with r3JUNV-GFP (Candid#1 strain) (55) and treated infected cells with C16 (3 µM) or C22 (3 µM) compounds. At the indicated hpi, TCS were collected, and viral titers were determined. Treatment with compound C16, but not with compound C22, caused 3 logs reduction in virus titers, at 48 and 72h pi, compared to vehicle control treated cells (Fig. 7A). Similarly, treatment with C16, but not with C22, inhibited propagation of LASV (Fig. 7B). As with LCMV (Fig. 1), genetic ablation of PKR resulted in enhanced propagation of LASV (Fig. 7 C).

**Figure 7.**
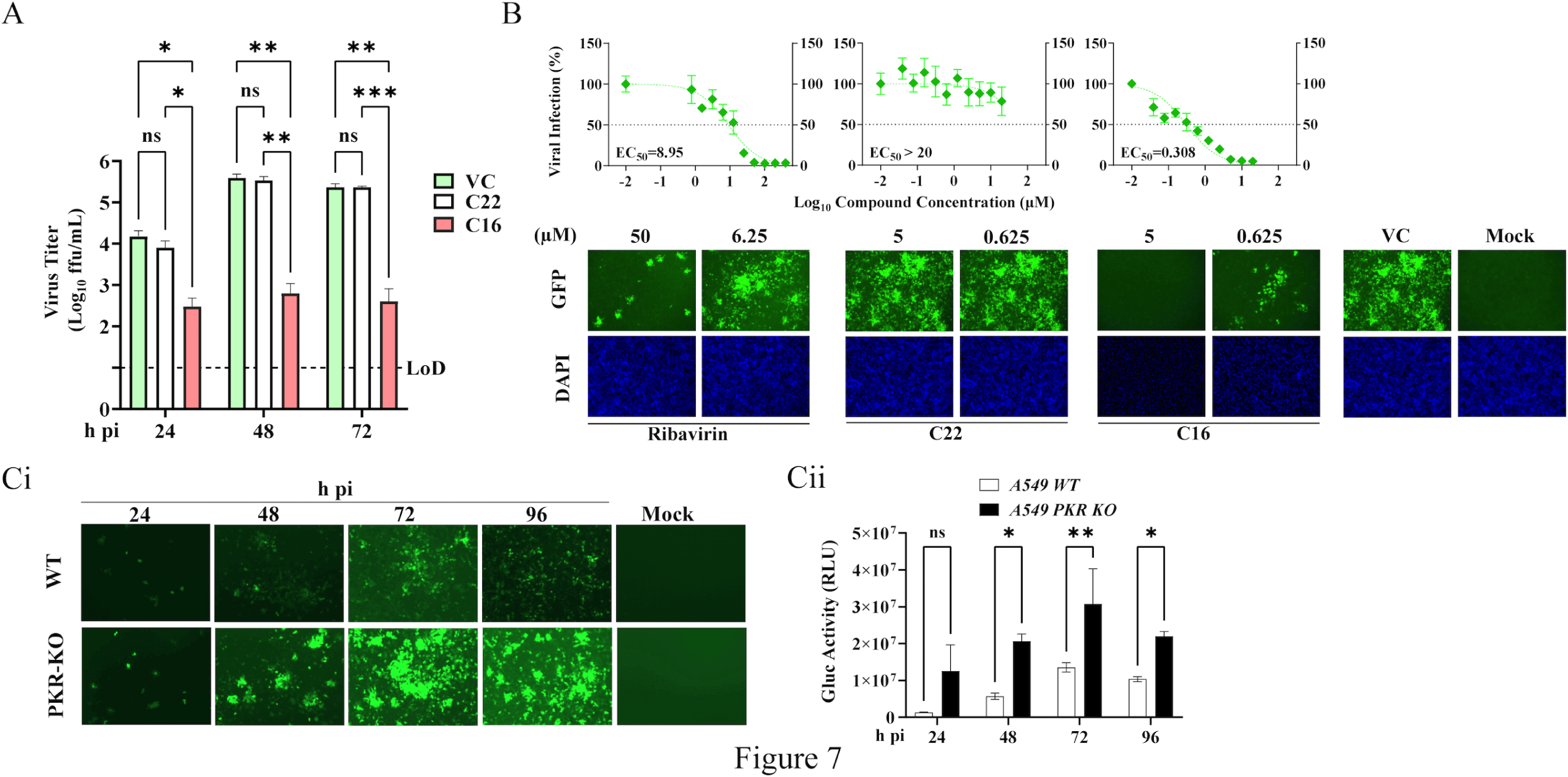
Contribution of PKR to JUNV and LASV multiplication in cultured cells. **A.** Treatment with compound C16 inhibits multiplication of JUNV. A549 cells were seeded into an M24 well plate (1.5x105 cells/well) and next day (18 h), they were infected with r3JUNV-GFP (Candid#1 strain) (55) at MOI of 0.05. After 90 minutes of adsorption (0.2 mL/well), the inoculum was removed, cells were washed once, and fresh medium (0.5 mL/well) containing C16 (3 µM) or C22 (3 µM) compounds, or VC, was added to the cells. At the indicated time points TCS samples were collected and virus titers were determined by FFUA on Vero E6 cells. **B.** The PKR inhibitor compound C16 exhibits a dose-dependent inhibitory effect on multiplication of LASV. A549 cells were infected with r3LASV (MOI 0.001). After 1 h of viral adsorption, compounds C16, and C22, or RBV were added at a range of concentrations. At 72 h post-infection, cell culture supernatants were collected and Gluc activity was quantified. Data were normalized by assigning the value of 100 to vehicle control (VC)-treated and r3LASV-infected cells. Cells were then fixed in 10% formalin and imaged for GFP expression using an EVOS M5000 imaging system. **C.** Genetic ablation of PKR results in enhanced r3LASV propagation. Parental and PKR KO A549 cells were infected with r3LASV- GFP/Gluc (MOI 0.001). At indicated time points post-infection, TCS samples were collected, and cells were fixed. GFP+ cells, corresponding to LASV-infected cells, were visualized by epifluorescence using an EVOS M5000 imaging (**Ci**). TCS samples were assayed for their Gluc activity (**Cii**). Two-way ANOVA with Šidák correction for multiple comparisons was implemented for statistical analysis.

## DISCUSSION

The PKR pathway is one of the major arms of cell innate immunity response to viral infections. Accordingly, viruses have evolved different strategies to counteract the PKR pathway. In this work, we have documented that LCMV/WT infection does not result in robust PKR activation, as determined by its autophosphorylation. This finding is consistent with those reported by others showing that at early times of infection and up to 48h pi, LCMV/WT-infected cells showed negligible activation of PKR (37), as well as results documented with other mammarenaviruses (41). Interestingly, LCMV multiplication was increased in PKR-KO cells, reflected in consistently 5 to 10-fold higher virus titers in PKR-KO than in WT A549 cells. It seems unlikely that LCMV actively interferes with PKR activation as A549-LCMV infected cells superinfected with SINV exhibited robust PKR activation. In contrast to LCMV/WT, infection of A549 cells with rLCMV/NP(D382A) resulted in robust PKR activation, which correlated with a severely restricted multiplication of rLCMV/NP(D382A) that was alleviated in PKR-KO A549 cells. PKR reconstitution studies using transfection of PKR-KO A549 cells with a PKR-expressing plasmid showed that multiplication of rLCMV/NP(D382) was restricted in PKR-KO A549 cells upon reconstitution of PKR expression, further confirming that PKR can exert an anti-LCMV activity. PKR was found to be involved in apoptosis, the formation of stress granules (SGs) (56, 57) and IFN-induced cellular necrosis (58).

However, neither total protein levels nor numbers of apoptotic cells were significantly affected in cells infected with rLCMV/NP(D382A) despite activation of eIF2α, a finding that warrants further studies.

PKR is activated by conformational change and autophosphorylation upon binding of dsRNA generated during viral genome replication. The 9D5 antibody has been reported to detect dsRNA species ≥ 40 bp (47, 48, 59) with a motif A_2_N_9_A_3_N_9_A_2_ (59), which is present in both GPC (802–826) and L (763-787, 1007-1031, 2970-2994, 3711-3735, 5974-5998) LCMV genes. However, our repeated attempts to detect dsRNA in A549 cells infected with LCMV/WT or rLCMV/NP(D382A) using the 9D5 antibody were unsuccessful, whereas we readily detected dsRNA in cells infected with SINV. Unexpectedly and in contrast with published findings (40), the 9D5 antibody was also unable to detect dsRNA in cells infected with JUNV. Nevertheless, our findings are consistent with earlier observations documenting the detection of dsRNA by the 9D5 antibody in cells infected with dsRNA and positive, but not negative, stranded RNA viruses (60). It is possible that virus-specific dsRNA species generated in LCMV infected cells are < 40 bp in length either because of a fast binding of NP to the newly formed RNA (61) or the lack of immediate supercoiling of RNA (62) which might prevent the formation of the structural binding site for the 9D5 antibody. Alternatively, it might be possible that the 9D5 antibody is targeting a specific domain that is common in dsRNA and +ssRNA supercoiled structures with sizes ≥ 40bp (62). The 9D5 antibody has been shown to have an affinity for the peptide motif SIGNAYSMFYDG (63), not present in LCMV proteins, and therefore it cannot be ruled out that the 9D5 antibody binds to a protein target expressed only under specific situations that were not recreated under our experimental conditions.

Our results with PKR deficient cells supported an expected antiviral activity of PKR, but intriguingly pharmacological inhibition of PKR activation uncovered a pro-viral activity of PKR. The PKR inhibitor C16 has been shown to affect PKR-independent biochemical intracellular transduction mechanisms (64), and the activity of cyclin-dependent kinases CDK2 and CDK5 (65), which could have contributed to the C16 antiviral activity against LCMV, LASV and JUNV. However, we have documented that siRNA-mediated knock-down of the CDK2 activator cyclin A2 protein resulted only in 50% reduction in levels of LCMV multiplication (66), which cannot account for the over 5 logs reduction in production of LCMV infectious progeny caused by C16 treatment we document in the present work. C16 has been extensively characterized for its inhibitory effect on autophosphorylation of PKR (49). In addition, comprehensive kinomics studies showed that C16 has high specificity for PKR with a Gini score of 0.44 PKR (67).

Moreover, other proposed kinase targets of C16 including JNK, MKKs, and GSK3β are downstream PKR pathway, which may reflect an upstream targeting pf PKR (68–70). Importantly, the C16 closely related compound C22 does not inhibit PKR autophosphorylation (49), which was associated with its lack of anti-mammarenavirus activity, providing additional evidence supporting inhibition of autophosphorylation of PKR as the main factor contributing to the C16 anti-mammarenaviral activity.

PKR may exert functions in LCMV-infected cells that do not require its activation. Thus, PKR has been shown to activate IKK by protein-protein interactions, leading to NF-kB activation and the induction of the T1IFN pathway (71). PKR has also been implicated in the inhibition of gelsolin-mediated regulation of actin dynamics and cytoskeletal cellular functions contributing to innate immunity including restricted virus cell entry, activities that do not appear to require PKR activation via autophosphorylation (72). The actin filament severing activity of gelsolin is inhibited by the oligomeric molecular chaperone CCT (73), and inhibition of the CCT activity resulted in restricted multiplication of LCMV (74), suggesting a connection between PKR, the host cell cytoskeletal activity, and mammarenavirus multiplication.

Similar to findings reported for infectious pancreatic necrosis virus (IPNV) (75), and SARS-CoV-2 (76) treatment with the PKR inhibitor C16 resulted in strong inhibition of LCMV multiplication in WT A549, supporting the role of PKR in LCMV multiplication, and uncovering PKR activation as a druggable target for the development of antiviral drugs against human pathogenic mammarenaviruses, including LCMV for which evidence indicates it is a neglected human pathogen of clinical significance (4–8), and a serious risk to immunocompromised individuals (9, 10). C16 has shown tolerability and efficacy in a mouse model of HCV-induced hepatocarcinogenesis (77), supporting its future evaluation, beyond the scope of the present work, in mouse models of LCMV infection.

## Materials and Methods

### Antibodies and compounds

We used the following antibodies at the indicated dilutions: Rat mAb VL4 to LCMV NP (BE0106, Bio X cells) 1 in 1000; mouse mAb 9D5 anti-dsRNA (3361, EMD Millipore) 1 in 2; mouse mAb rJ2 (MABE1134, MILLIPORE/SIGMA) 1 in 125, rabbit mAb anti-PKR (D7F7) (12297S, Cell signaling technology) 1 in 1000; antibody [E120] anti-pPKR (phosphor T446) (ab32036, Abcam) 1 in 1000; rabbit mAb anti-peIF2α (Ser51) (D9G8) XP® (3398T, Cell Signaling); Phospho- eIF2α-S51 mouse mAb (AP0692, Abclonal) 1:1000; mouse Anti-LCMV-GP hybridoma G204 1 in 500, goat anti-rabbit IgG (H+L) cross-adsorbed secondary antibody, Alexa Fluor 488 (A-11008, Thermo Fisher Scientific); goat anti-mouse secondary antibody, Alexa Fluor 568 (Thermo Fisher Scientific). PKR Inhibitor C16 (15323-1, Thermo Fisher Scientific), PKR Inhibitor, Negative Control (C22) (52745510MG, Fisher Scientific).

### Plasmids

pSB819-PKR-hum was a gift from Harmit Malik (Addgene plasmid # 20030; http://n2t.net/addgene:20030 ; RRID:Addgene_20030) (78) and it was used to subclone the human PKR isoform 2 into pCAGGS plasmid preceded by mCherry and separated by P2A self-cleaving peptide sequence (43). The PKR was HA tagged at its C-terminus. HD In-Fusion kit (Takara 638909 In-Fusion® HD Cloning Plus) was used for cloning.

### Cell lines

A549 (*Homo sapiens)* (American Type Culture Collection, ATCC, CCL-185), A549 PKR-knockout, A549 RNase L-knockout, A549 MAVS-knockout (79), BHK-21 (*Mesocricetus auratus*) (ATCC, CCL-10), and Vero E6 (*Chlorocebus aethiops*) (ATCC CRL-1586) cell lines were maintained in Dulbecco’s modified Eagle’s medium (DMEM) (Thermo Fisher Scientific) supplemented with 10% heat-inactivated fetal bovine serum (FBS), 2 mM L-glutamine, 100 µg/ml of streptomycin, and 100 U/ml of penicillin. BHK-21 medium also contained 5% tryptose phosphate broth.

### Viruses

The recombinant LCMV, clone 13 (Cl-13) has been described (80), rLCMV/NP(D382A) was rescued via reverse genetics as described (80). Mutation D382A in NP was generated with the In-Fusion cloning method (Takara 638909 In- Fusion® HD Cloning Plus). rLCMV/NP(D382A) was grown in a BHK-21 cells and its identity confirmed by sequencing. rSINV-GFP was obtained from Matthew Daugherty (UCSD). The recombinant trisegmented (r3) LASV, strain Josiah, expressing GFP and Gluc was rescued as described for other r3MaAv, including LCMV (81), JUNV (82), and TCRV (83), using described LASV reverse genetic approaches (84, 85). Briefly, the mPolI-S LASV plasmid was modified to contain two restrictions sites (BsmBI and BbsI) in opposite orientation (mPolI-S LASV backbone). To generate the mPolI-S1 LASV, the open reading frame (ORF) of EGFP was amplified by PCR and cloned into the BsmBI restriction site of the mPolI-S LASV backbone. Then, the ORF of LASV GPC was amplified by PCR and cloned using BbsI. To generate the mPolI-S2 LASV, the ORF of LASV NP was amplified and cloned into the BsmBI restriction site of the mPolI-S LASV backbone. Then, the ORF of Gluc was amplified and cloned using BbsI. All the plasmids were entirely sequenced by Plasmidsaurus to ensure the absence of unwanted mutations. To generate r3LASV, BHK21 cells in 6-well plates were transfected with mPolI-L LASV, mPol-S1 LASV, and mPol-S2 LASV, using LPF2000. At 48 h post-transfection, BHK21 cells were scaled up into a T75 flask, and 48 h after the presence of the r3LASV was verified by GFP expression. Cell culture supernatants were collected to confirm Gluc expression and to generate viral stocks. Stocks of r3LASV were generated and titrated in Vero E6 cells (84, 85). All the experiments with r3LASV were conducted in the biosafety level 4 (BSL4) laboratory at Texas Biomedical Research Institute. Protocols were approved by Texas Biomedical Research Institute Biosafety (BSC) and Recombinant DNA (RDC) committees (BSC21-013 and RDC21-013, respectively).

### Virus titration

Virus titers were determined by focus forming assay (FFA) using Vero E6 cells. Briefly, cells were seeded in 96-well plates at a density of 2 × 10^4^ cells/well and fixed at 20 h pi using 4% paraformaldehyde in phosphate-buffered saline (PBS) for 25 minutes, washed twice with PBS, permeabilized with 0.3% Triton X-100 containing 3% BSA in PBS, incubated with the rat mAb VL4 against LCMV NP (Bio X Cell) for 1 h at room temperature, and then incubated with an anti-rat IgG conjugated to Alexa Fluor 568 for 1 h, cells were washed 3X with PBS after each antibody probing step prior to imaging.

### RT-qPCR

Cells were infected with LCMV (MOI 0.01) or rLCMV/NP(D382A) (MOI 0.05). Total cellular RNA was isolated using TRI reagent (TR 118) (MRC), and 1 µg reverse-transcribed to cDNA using the SuperScript^TM^ IV first-strand synthesis system (Thermo Fisher Scientific). Powerup SYBR (A25742, Life Technologies) was used to amplify LCMV NP and the housekeeping gene GAPDH using the following primers: NP forward (F): 5′ CAGAAATGTTGATGCTGGACTGC-3′, and NP reverse (R): 5′- CAGACCTTGGCTTGCTTTACACAG-3′ (86); GAPDH F: 5′- CATGAGAAGTATGACAACAGCC-3′, and GAPDH R: 5′-TGAGTCCTTCCACGATACC- 3′. ISG15 F: 5′-CAGGACGACCTGTTCTGGC-3′, and ISG15 R: 5′-GATTCATGAACACGGTGCTCAGG-3′, MX1 F: 5′- GCAGCTTCAGAAGGCCATGC-3′, MX1 R: 5′-CCTTCAGGAACTTCCGCTTGTC-3′.

### Western blotting

Cell monolayers were washed with ice-cold PBS and whole cell lysates prepared in cytoplasmic lysis buffer (50mM Tris HCl, 150mM NaCl, 1% NP-40, 10% glycerol, 2mM EDTA) supplemented with Halt Protease and Phosphatase Inhibitor Cocktails (PI78442, Thermo Fisher Scientific). Lysates were clarified by centrifugation at 10,000 RCF for 20 minutes. Samples (24 µg) were denatured by heating 5 minutes at 95°C and separated by Stain-Free SDS-PAGE gel (4568096, Bio-Rad). The gel was activated using one-minute setting (Bio-Rad Imager, imaged, then transferred to the low fluorescence polyvinylidene difluoride (PVDF) membrane (1704274, Bio-Rad) that was probed with mAbs to PKR (Cell Signaling Technology) or phospho-PKR-446 (AP1134, Abclonal). Following the incubation with a secondary HRP antibody, the bands were visualized with the chemiluminescent substrate (Thermo Fisher Scientific). The bands’ signals were quantified and analyzed using Image Lab V6 (Bio-Rad).

### Northern blotting

RNA samples (5 µg) were fractionated by 2.2 M formaldehyde-agarose (1.2%) gel electrophoresis. The gel was washed once with warm each H_2_O and 10 mM NaPO_4_, and RNA transferred in 20 x SSC [3 M sodium chloride, 0.3 M sodium citrate] to a Magnagraph membrane (NJTHYA0010, Osmonics MagnaGraph nylon) using the rapid downward transfer system (TurboBlotter). Membrane-bound RNA was cross-linked by exposure to UV light, the membrane was washed with MilliQ water and stained with methylene blue (MB) to reveal the 18S and 28S RNA plus RNA ladder.

After photo documentation, 1% SDS solution was used to remove MB staining, and the membrane was hybridized using QuickHyb (#201220-12 Agilent) to a ^32^P-labeled dsDNA NP. Hybridization was performed at 65°C overnight The DNA probe was prepared according to the supplier’s protocol using a DecaPrime kit (Ambion). After overnight hybridization, the membrane was washed twice with 2 x SSC–0.2% SDS at 65°C, followed by two washes with 0.2× SSC–0.2% SDS at 65°C, and then exposed to an X-ray film.

### Immunofluorescence

Individual images were obtained using the Keyence BZ-X710. Files containing the labeled images were transferred to a laptop for processing the data using ImageJ. PowerPoint (2019 version) was used to compile and arrange the individual images. Each image was imported individually and organized within the corresponding composite, adjusting the canvas size to ensure a cohesive layout. Adobe Illustrator was used to align the panels within the composite.

### Dose-Response and Cell viability assay

CellTiter 96 AQueous One Solution (G3580, Promega) was used, according to manufacturer protocol, to quantify viable cells. DAPI and GFP signals were quantified after fixation of cells with 4% PFA using a Biotek H4 plate reader, and the Celigo Image Cytometer (Nexcelom). Data were normalized using vehicle-treated infected and mock-infected cells as the highest and lowest, respectively, signals. Ribavirin (100μM) was used as a control of a validated inhibitor of LCMV multiplication. Four replicates were used to quantify each sample.

### Flow Cytometry

Cells were washed and treated with Accutase (490007-741, VWR), transferred into 15 mL tubes, washed twice with FACS buffer (1X PBS, 2% FBS, 1mM EDTA), then fixed with 4% PFA for 20 minutes, transferred to a 96 well plate (round bottom). After permeabilization for 30 minutes (Permeabilization Buffer, 00-8333-56, eBioscience), cells were washed and reacted with the indicated antibodies. Cells were washed twice with permeabilization buffer followed by centrifugation for 3 minutes after each antibody probe. The primary antibody was probed for 1 h, secondary antibody for 30 minutes. Cells were resuspended in FACS buffer for analysis.

### Dose-response inhibition of r3LASV

A549 cells (96-welll plate format, quadruplicates) were infected with r3LASV (MOI 0.001). After 1 h of viral adsorption, indicated concentrations of inhibitors were added. At 72 h post-infection, cell culture supernatants were collected and Gluc activity was quantified according to the manufacturer protocol using Pierce™ Gaussia Luciferase Glow Assay Kit (16160, Thermo Fisher Scientific) and GloMax plate reader (Promega). Data were normalized to mock-treated, r3LASV-infected cells. Cells were then fixed in 10% formalin and imaged for GFP expression using an EVOS M5000 imaging system (Thermo Fisher Scientific).

### Infection of parental and PKR KO A549 cells with r3LASV-GFP

Parental and PKR KO A549 cells (96-welll plate format, triplicates) were infected with r3LASV-GFP/Gluc (MOI 0.001). At indicated timepoints post-infection, cell culture supernatants were collected and Gluc activity was quantified according to the manufacturer protocol using Pierce™ Gaussia Luciferase Glow Assay Kit (16160, Thermo Fisher Scientific) and GloMax plate reader (Promega). Data were normalized by subtracting Gluc signal from mock-infected cells. Cells were then fixed in 10% formalin and imaged for GFP using an EVOS M5000 imaging system.

### Statistical analysis

Differences that are statistically significant in virus growth and NP gene expression were determined using an unpaired t-test with Welch correction.

Bands’ signals of IB were quantified using Image Lab (Bio-Rad). All statistical analyses were conducted using GraphPad Prism software v9.5.1 (GraphPad). Flow cytometry analyses were performed using FlowJo v10.9 software.

## Acknowledgments

This research was supported by NIH/NIAID grant RO1 AI142985 to JCT and NIH T32 AI007354 (ASK). ASK was supported in part by Open Philanthropy and the Life Sciences Research Foundation. This is manuscript 30241 from The Scripps Research Institute. We want to thank Olena Shtanko for her help and assistance with the LASV experiments at BSL4.

